# Integrative analysis of Zika virus genome RNA structure reveals critical determinants of viral infectivity

**DOI:** 10.1101/412577

**Authors:** Pan Li, Yifan Wei, Miao Mei, Lei Tang, Lei Sun, Wenze Huang, Jianyu Zhou, Chunlin Zou, Shaojun Zhang, Cheng-feng Qin, Tao Jiang, Jianfeng Dai, Xu Tan, Qiangfeng Cliff Zhang

**Author notes:** Co-first author (Pan Li, Yifan Wei, Miao Mei, Lei Tang). Correspondence (X.T.), (Q.C.Z.).

## Abstract

Since its outbreak in 2007, Zika virus (ZIKV) has become a global health threat that causes severe neurological conditions. Here we perform a comparative *in vivo* structural analysis of the RNA genomes of two ZIKV strains to decipher the regulation of their infection at the RNA level. Our analysis identified both known and novel functional RNA structural elements. We discovered a functional long-range intramolecular interaction specific for the Asian epidemic strains, which contributes to their infectivity. Our findings illuminate the structural basis of ZIKV regulation and provide a rich resource for the discovery of RNA structural elements that are important for ZIKV infection.

## INTRODUCTION

Zika virus (ZIKV) is a mosquito-borne single-stranded RNA virus of the *flaviviridae* family (Petersen et al., 2016). The virus was first isolated in Uganda in 1947 and had remained obscure until its outbreak in Micronesia in 2007. Since then it has caused epidemics in Pacific islands and later in America and Asia. In response to its rapid spread and association with serious brain disorders, including microcephaly in newborns and Guillain-Barre syndrome in adults, the World Health Organization has declared it a public health emergency and tremendous efforts have been devoted to understanding its pathogenesis and to developing effective treatments (Brasil et al., 2016, Cugola et al., 2016, Mlakar et al., 2016). However, there remains no specific therapy or approved vaccine for ZIKV infection.

The ZIKV genome is an approximately 10.8 kb positive-sense RNA in an mRNA-like pattern (Zhu et al., 2015). ZIKV strains can be classified into the ancestral African lineage and the contemporary Asian lineage based on genome sequences, with the latter responsible for the current epidemics. To understand how the Asian strains lead to epidemics and illness, a crucial step is to identify genomic variations that affect their infectivity and pathogenicity. Comparative studies have focused on mutations that change protein sequences. For example, it has been proposed that a key amino acid substitution S139N in the prM protein contributes to fetal microcephaly (Yuan et al., 2017), and another mutation A188V in the NS1 protein promotes transmissibility in mosquito vectors (Liu et al., 2017). However, most genomic variations between the two lineages are synonymous or non-coding, and thus are not expected to affect the encoded proteins. We analyzed a set of representative ZIKV strains and found on average 1,166 mutations between the Asian strains and the African strains. Among them, about 88%, i.e., 1,020 mutations, are synonymous or in non-coding UTR regions (Supplementary Figure S1A, S1B and S1C). Whether and how these mutations could also contribute to ZIKV infection remain to be answered.

It is well-known that the genomic RNA of flaviviruses participates in viral processes including translation, replication, packaging and evasion of host cell antiviral responses (Rodenhuis-Zybert et al., 2010). RNA structural elements are functionally involved in many of these processes. For example, conserved multi-pseudoknot structures in the 3′ UTR of ZIKV and other flaviviruses can stall the RNA exonuclease Xrn1, thereby giving rise to sub-genomic flavivirus RNAs that help the virus evade cellular antiviral processes (Akiyama et al., 2016, Filomatori et al., 2017). In addition, an intramolecular RNA-RNA pairing between the 5′- and the 3′-UTRs facilitates transformation between the linear and circular conformations of the genomic RNA, and thus plays an important in role coordinating virus replication (Villordo et al., 2009). To date, our knowledge of ZIKV RNA structure is mainly limited to the untranslated regions. However, the coding region constitutes over 95% of the ZIKV genome and likely contains a wealth of functional structural elements yet to be discovered. Uncovering these structural elements and their differences between the two lineages may reveal a molecular rationale for the outbreak of epidemics.

Next-generation sequencing-based technologies enable the profiling of the complete genome structures of RNA viruses. A pioneer study of the HIV RNA genome structure discovered known and unrecognized regulatory motifs, as well as a higher-order organizational principle that RNA structures directly encode protein domain junctions (Watts et al., 2009). Other studies examined multiple HCV genome structures and found various structural regulatory elements across the whole genome including coding regions (Mauger et al., 2015, Pirakitikulr et al., 2016). The studies suggested that the viral RNA structures may have evolved into a sophisticated complex network that protects the genome from both RNase L and double-stranded RNA-induced innate immune sensors. And conformational changes within these structural motifs can influence viral replication or immune evasion. However, all these studies are based on *in vitro* experiments.

Here we investigated RNA secondary structures and intramolecular RNA-RNA interactions of two ZIKV RNA genomes *in vivo* by combing two orthogonal high throughput sequencing based technologies, icSHAPE (Spitale et al., 2015) and PARIS (Lu et al., 2016) (Figure 1). We generated maps of the secondary structures of two ZIKV strains representing the Asian and African lineages, with many novel long-range intramolecular RNA-RNA interactions including common and lineage-specific structural elements. We identified a functionally important long-range intramolecular interaction that is specific for the Asian strains after 2007 and regulates their infection in certain cell lines. Therefore, we provide a rich resource for understanding the structure-function relationship of ZIKV genomic RNA.

**Figure 1.**
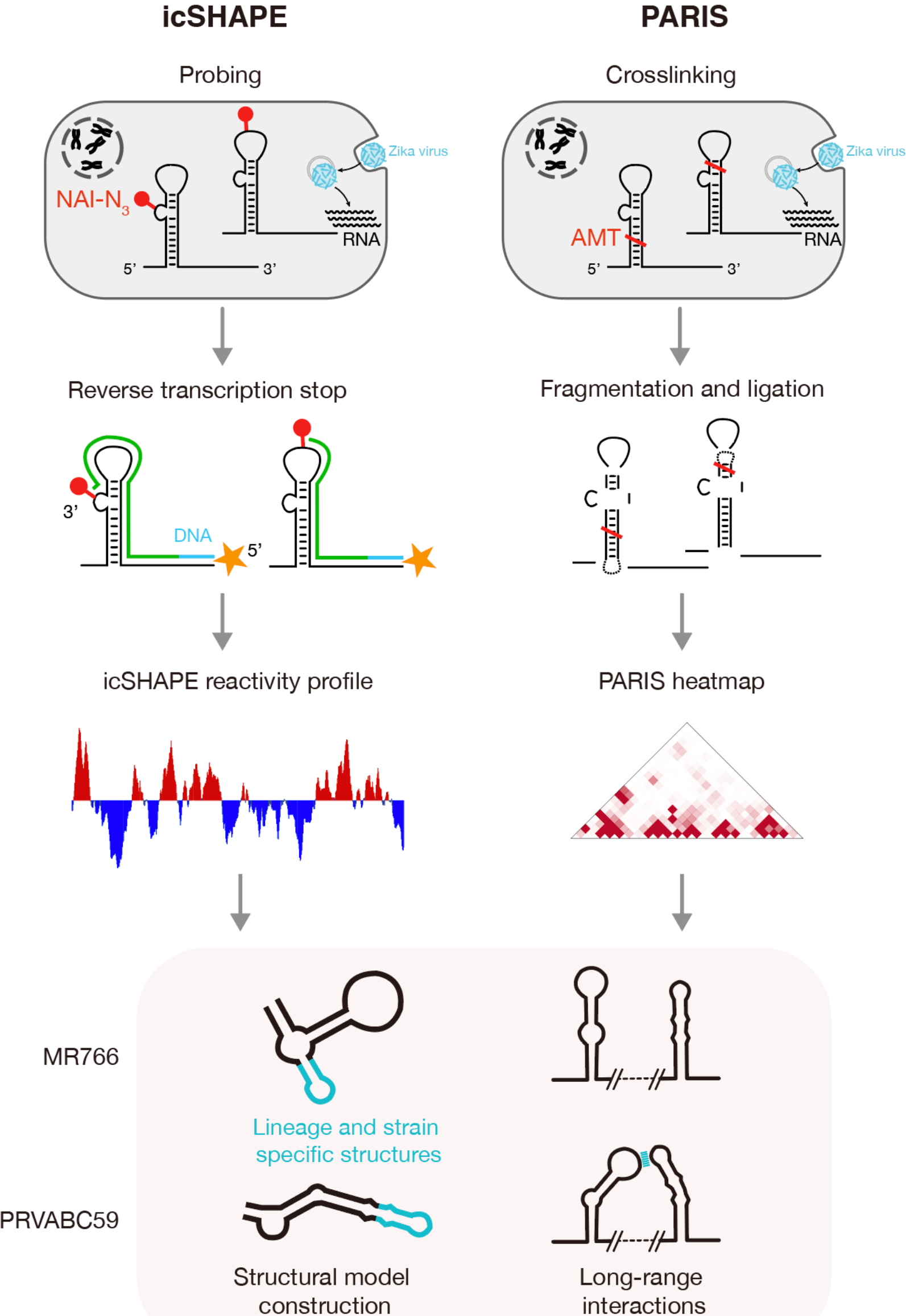
Schematic presentation of ZIKV RNA structure *in vivo* study by the combination of icSHAPE and PARIS. See also STAR Methods, Figure S1. For ZIKV PRVABC59 and MR766 h.p.i Huh-7 cells, icSHAPE is performed to detect virus RNA flexibility (paired or unpaired) at nucleotide resolution *in vivo*. PARIS was performed to detect intramolecular interactions of ZIKV RNA genomes *in vivo*.

## RESULTS

### icSHAPE defines nucleotide flexibility profiles of ZIKV genomes

For RNA structural determination, we chose PRVABC59 (Lanciotti et al., 2016) as a representative Asian strain and MR766 (Dick et al., 1952) for the African strains, according to their positions in the ZIKV phylogenetic tree (Supplementary Figure S1A). We noticed that the PRVABC59 reference genome from GenBank lacks a conserved functional small hairpin in the 3′UTR, so we re-assembled the genomes with our sequencing data and completed the sequences. The two ZIKV genomes displayed 6.5% and 3.9% sequence variation within the 5′- and 3′-UTR regions, respectively, and 11.3% variation within the polyprotein coding region (Supplementary Figure S1D). The elevated sequence conservation within the UTRs relative to the coding regions reflects the importance of known functional RNA elements in flavivirus UTRs.

First, we performed “*in vivo* click selective 2-hydroxyl acylation and profiling experiments” (icSHAPE) to measure the structural flexibility of every nucleotide within the PRVABC59 and MR766 RNA genomes in infected cells (Figure 2). We treated ZIKV-infected Huh7 cells with the icSHAPE reagent NAI-N3, which preferentially reacts with unstructured and flexible nucleotides. We then purified the modified RNA and performed reverse transcription. NAI-N3-modified bases block reverse transcriptase, yielding cDNA fragments that end at the modified site. We performed deep sequencing and computational analyses to map the reverse transcription termination sites, generating a flexibility or icSHAPE reactivity score for each nucleotide. Flexibility negatively correlates with the likelihood of secondary structure, thus providing a measure of the pairing probability of each nucleotide

**Figure 2.**
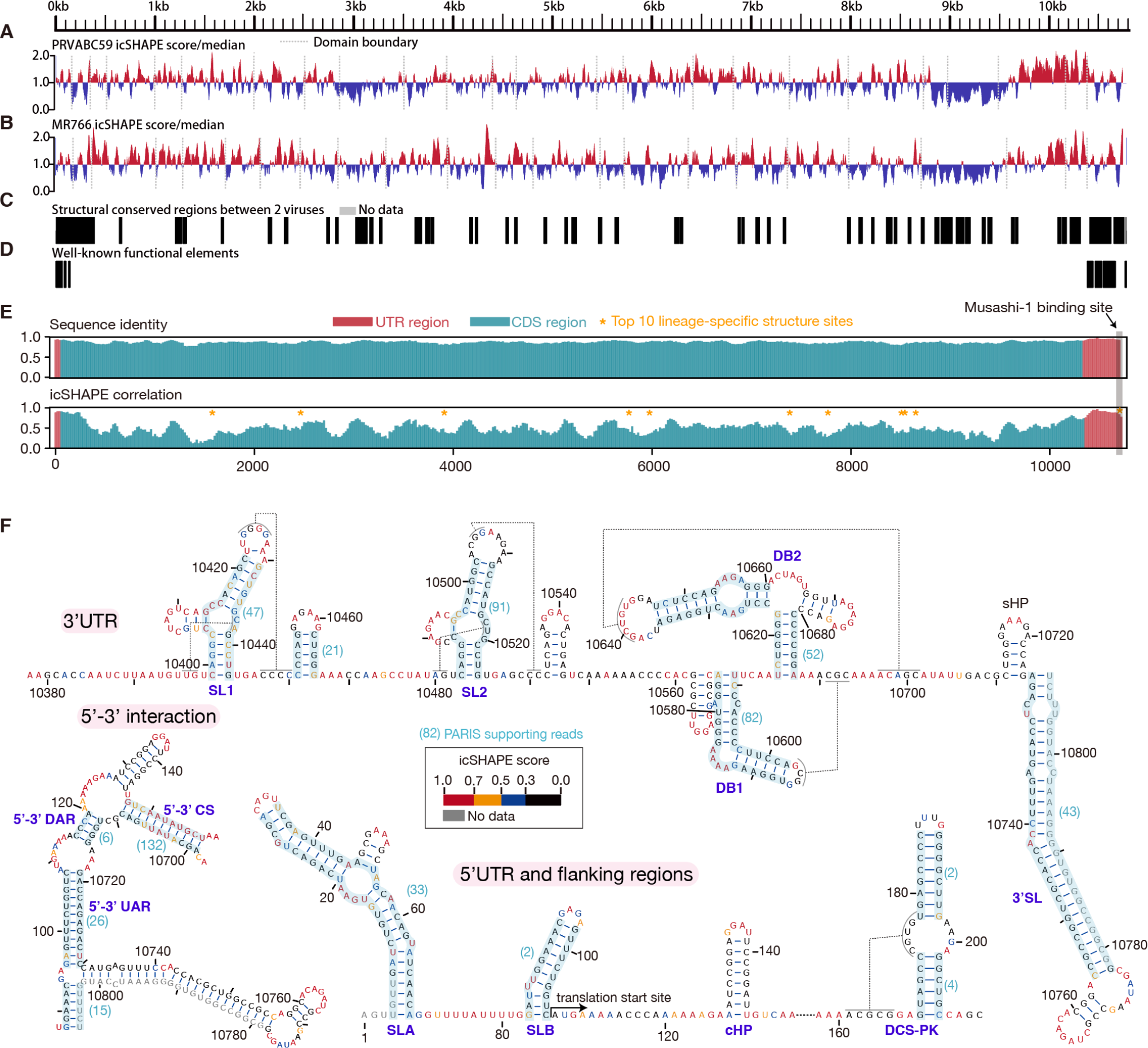
Structural overview of icSHAPE profiling in the genome of ZIKV PRVABC59 and MR766. See also STAR Methods, Tables S1 and S3. A. Normalized *in vivo* icSHAPE reactivity score of PRVABC59 shown relative to the global median value, with higher values correspond to more flexible nucleotides. Blue color thus represents a region more likely to be pairing state, and red color represents more likely a non-pairing region. Normalized scores are smoothed with a 30-nt window. B. Normalized *in vivo* icSHAPE reactivity score of MR766. C. Structure-conserved regions between two viruses, and it is defined as those regions with icSHAPE difference more than 0.15 in a 30-nt window. D. Well-known functional elements in two viruses as showed in F. E. Sequence identity and correlations of icSHAPE reactivity scores between PRVABC59 and MR766 strains in sliding windows. The CDS region and UTRs are respectively colored with cyan and red. Sites marked with yellow asterisk are top-10 lineage-specific structure sites between PRVABC59 lineage and MR766 lineage. F. Known structural models of the 3′UTR (top), the 5′UTR and flanking regions (bottom middle) and the 5′-3′ interaction (bottom left), colored with icSHAPE reactivity scores. Dashed lines represent pseudoknots. RNA-RNA interactions with PARIS data supporting are shadowed in light blue with gapped reads numbers in brackets.

We obtained icSHAPE reactivity scores for over 99.6% of the nucleotides within the two ZIKV genomes (Figure 2A and 2B, Supplementary Table S1). The icSHAPE scores were reproducible between independent biological replicates (R ≥ 0.89). The structural conservation was significant and contains many well-known functional elements in flaviviruses (Figure 2C and 2D). Consistent with previous findings, we noticed that the structural conservation by icSHAPE scores was lower than the degree of the sequence conservation between the two strains (R=0.52; Figure 2E). Five RNA elements in the 3′-UTR, including Stem-loop1 (SL1), Stem-loop2 (SL2), Dumbbell1 or pseudo-Dumbbell (DB1), Dumbbell2 (DB2) and the small hairpin 3′ Stem-loop (sHP-3′SL), are conserved structures and established benchmarks for RNA folding prediction (10). All these elements, except sHP-3′SL, are at the 5′ end of sub-genomic flavivirus RNAs and may function as Xrn1 RNase resistant components in ZIKV (9).

At the 5′-end of the viruses, three stem loops, including stem loop A (SLA), stem loop B (SLB) and capsid hairpin (cHP), are defined from previous SHAPE and RNase digestion studies in Dengue virus (Lodeiro et al., 2009). The DCS-PK element in the 5′ UTR flanking region, is a novel pseudoknot found in coding region (Liu et al., 2013). SLA has been proposed to function in viral RNA synthesis, whereas SLB and cHP form base pairs with the 3′ end for genome cyclization and the DCS-PK element helps enhance genome cyclization during replication (Liu et al., 2013). Generally, unpaired nucleotides within these structural elements had higher icSHAPE scores, whereas paired nucleotides had lower icSHAPE scores, demonstrating that icSHAPE accurately measured ZIKV genomic RNA structures in infected cells (Area Under Curve, AUC=0.83, Figure 2F). The known long-range interaction between the 5′UTR and 3′UTR was also generally consistent with the icSHAPE scores. As one exception, the small stem of 5′-3′ CS was not reflected in the icSHAPE scores, which might indicate that this structure is dynamic or does not form under these conditions.

We also used icSHAPE to measure *in vitro* ZIKV RNA structures. RNA was extracted from ZIKV-infected Huh7 cells, refolded in an icSHAPE modification buffer for RNA structure stability, and then modified with NAI-N3 (Spitale et al., 2015). The remaining steps and data analysis were the same as for the *in vivo* icSHAPE measurement. We observed moderate correlations between *in vivo* and *in vitro* viral RNA structures (R=0.75 for both PRVABC59 and MR766), similar to that of eukaryote RNAs in previous studies (Spitale et al., 2015). Our data confirmed that RNA structures *in vivo* are generally more open than *in vitro*. Our *in vitro* icSHAPE data displayed lower agreement with the canonical 3′UTR RNA structure models than the *in vivo* data (AUC=0.74 vs. AUC=0.83), indicating that previous studies have recovered the *in vivo* conformations of many functional structural elements. Importantly, the difference between our *in vitro* and *in vivo* results implies that viral RNAs adopt distinct conformations in infected cells, highlighting the importance of studying RNA structures in their cellular context to uncover biologically relevant conformations.

### PARIS uncovers RNA-RNA interactions within ZIKV genomes

The RNA structural flexibility measured by icSHAPE scores represents the probability that a nucleotide is in a non-pairing, or single-stranded form. To directly map the RNA-RNA parings in ZIKV genomes, we used the newly developed PARIS method (Psoralen Analysis of RNA Interactions and Structures, Figure 1), which globally determines RNA duplex structures in live cells via reversible psoralen crosslinking (Lu et al., 2016). ZIKV-infected Huh7 cells were treated with the psoralen derivative AMT, and crosslinked RNA duplexes were purified by 2-D gel separation, proximity ligated and resolved by sequencing and bioinformatics analysis. An RNA-RNA interaction is defined by an alignment of gapped reads that can be mapped to the two stems of the duplex. Although PARIS cannot technically distinguish whether an RNA-RNA interaction is intramolecular or intermolecular, because proximity ligation is a key step of PARIS, and the interactions detected are typically intramolecular.

We discovered a large number of RNA-RNA interactions or pairing structures for both viruses by PARIS (Figure 3A and Supplementary Figure S2A). The PARIS data were reproducible between independent biological replicates (R=0.98), clustered into 1482 duplexes for PRVABC59 and 1282 duplexes for MR766 (Supplementary Table S2). Among these duplexes, we defined those with pairing distance of more than 1kb as long-range interactions, resulting in 230 for PRVABC59 (16% of all duplexes) and 178 for MR766 (14% of all duplexes) (Supplementary Table S2). Among the 15 short-range interactions of known local structures in the 3′UTR and 5′UTR with flanking regions, 10 were captured by PARIS, with the number of supporting reads ranging from 2 to 91 (data from the PRVABC59 experiment, Figure 2F). Only a few long-range interactions have been previously reported; among them is the interaction between the 5′UTR and the 3′UTR, which is highly conserved in flaviviruses and essential for their genome cyclization and replication (Liu et al., 2016). These interactions, including 5′-3′UAR, 5′-3′DAR and 5′-3′CS, are well-recovered in our results (Figure 2F).

**Figure 3.**
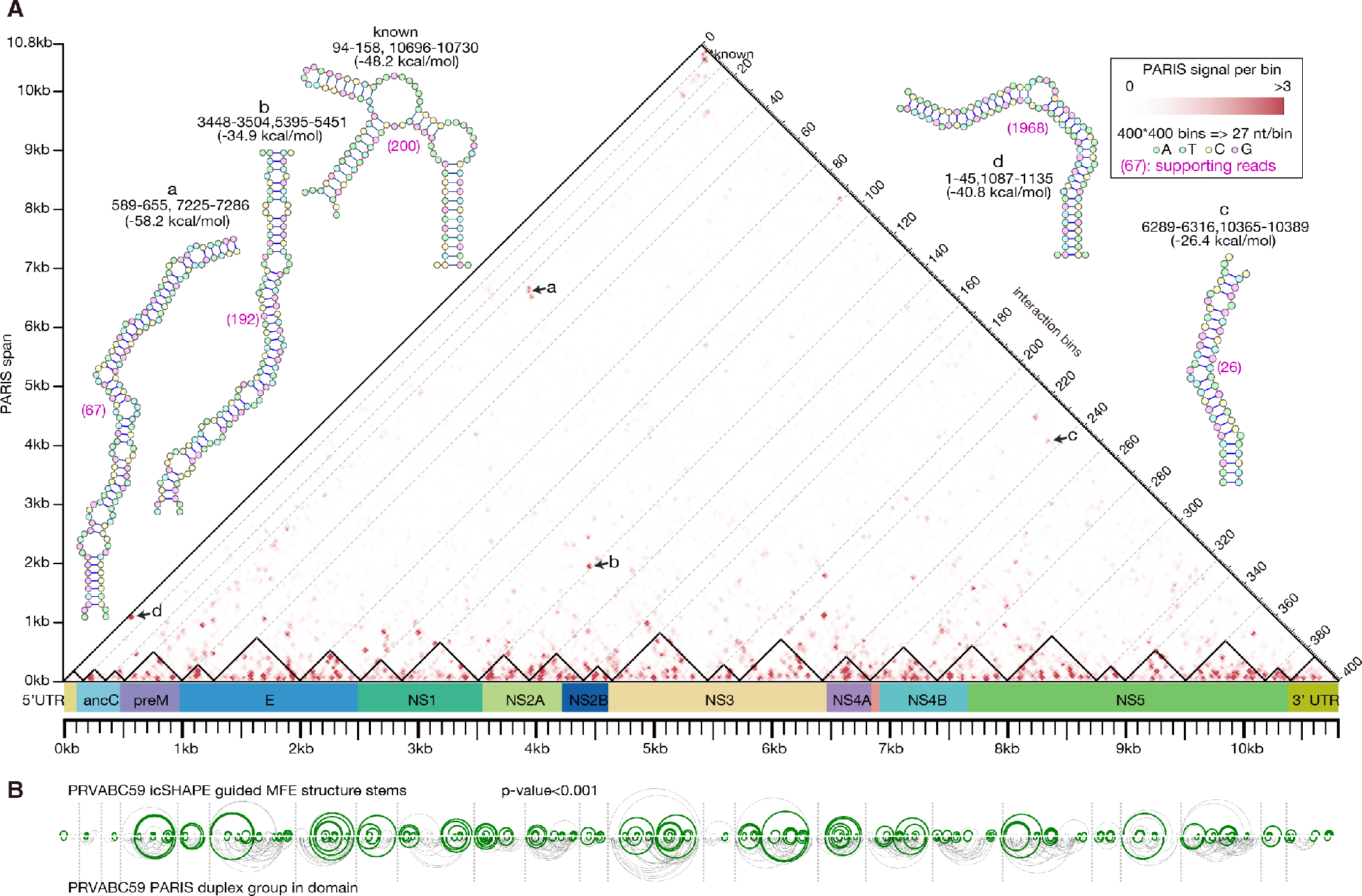
Structural overview of PARIS data in PRVABC59. See also STAR Methods, Figures S2, S3 and S4, and Tables S1 and S2. A. Heatmap of the PARIS connecting reads in the PRVABC59 genomes. Each data point represents an interaction between two regions. The coordinates of the two regions should be read by projecting the data point along the two declining axes. PARIS raw reads are normalized to signal which is showed as the color depth. Pairing structural models of some most stable long-range interactions including the known 5′-3′ interaction are highlighted with coordinates and folding energies (**a**~**d** and **known**). Small triangles flanking the middle bar represent RNA structural domains of PRVABC59. B. Comparison between icSHAPE-guided predicted base-pairing interactions by MFE (up) and PARIS interactions (below) for each domain of PRVABC59. Green arcs represent the common interactions while gray arcs are interactions only inferred from one method. P-value is calculated by shuffle predicted stems for 1000 times.

The single-strandedness measured by icSHAPE and the intramolecular interactions measured by PARIS provide two complementary sets of information on RNA structures. We noted that icSHAPE scores are usually very low for PARIS local structures, suggesting good agreement between the data. In addition, we used the software suite *RNAstructure* (Deigan et al., 2009) to predict a local minimum free-energy (MFE) structure with icSHAPE scores as a constraint (Figure 3B and Supplementary Figure S2B). We compared the base-pairing in the icSHAPE-assisted predicted structure with those measured by PARIS and found substantial overlap (p-value<0.001, permutation test, Figure 3B and Supplementary Figure S2B). However, it is worth noting that the agreement between icSHAPE and PARIS is lower for long-range interactions, which may reflect that many long-range interactions are transient, switching their conformations to open states under different conditions.

We noticed that short-range interactions are more conserved than long-range ones between the two ZIKV strains (Supplementary Figure S2C). We found that MR766 and PRVABC59 share over one third of short-range interactions, but less than 10% of long-range interactions. PARIS relies on psoralen intercalating in RNA helices, and requires deep sequencing due to limited crosslinking and ligation efficiencies (Lu et al., 2016). Our approach might underestimate the conservation of long-range interactions due to the limited coverage of PARIS, but this substantial difference still suggests lower evolutionary constraints for long-range than for short-range interactions.

To evaluate the plausibility of the detected long-range interactions, we analyzed their pairing energy. We first carried out a permutation test to determine the likelihood of an RNA pairing structure. For every PARIS long-range interaction, we calculated the pairing energy, using the *bifold* program from software suite *RNAstructure* with default parameters (Low et al., 2010, DiChiacchio et al., 2016). Next, we permutated the sequence in both stems 100 times, calculated the minimal fold free energy each time, and compared the true pairing energy with the averaged energy of 100 permutated ones. We found that most of the PARIS interactions are more stable than the permuted ones (p-value=6.2E-64 for PRVABC59 and p-value=1.5E-45 for MR766, pairwise t-test. Supplementary Figure S3A). This analysis demonstrates the specificity of PARIS experiments.

We also compared the folding energies of the long-range interactions discovered by PARIS with putative interactions between homologous regions of the other virus strain. Briefly, for each stem of a duplex defined by PARIS in one strain, we located their homologous regions in the other strain, and defined a putative homologous interaction by pairing the two homologous regions. We calculated the corresponding folding energies and found that most of the strain-specific PARIS interactions are much more stable than putative homologous ones (p-value=7.9E-24 for PRVABC59-specific long-range interactions, and p-value=2.8E-18 for MR766-specific ones, pairwise t-test. Supplementary Figure S3B), whereas the common interactions displayed similar folding energies (p-value=0.64, pairwise t-test, Supplementary Figure S3B). Together, our analyses suggest that many of the PARIS long-range interactions are energetically favorable. We highlighted some of the most stable long-range interactions in Figure 3A and Supplementary Figure S2A (labeled as **a**~**d** and **known**), including the well-known 5′-3′CS interaction. We hypothesize that some of the other long-range interactions are also important in virus infection.

We also used PARIS to measure ZIKV RNA-RNA interactions that form *in vitro*. We observed a lower fraction of long-range interactions in ZIKV folded *in vitro* compared to *in vivo* (p-value=1.68E-81 for PRVABC59, and p-value=1.50E-23 for MR766. Supplementary Figure S3C and S3D). This indicates the ZIKV is more extended in cells that forms less compacted short-range interactions.

### Architecture and structural models of ZIKV genomes

Interestingly, we observed that the intra-molecular interactions form clusters in the PARIS data heatmap (Figure 3A and Supplementary Figure S2A), similar to clusters found in genome connectivity maps from Hi-C data. This inspired us to define structural domains in each of the two ZIKV genomes. We implemented an algorithm that identifies clusters with dense intra-domain interactions. The algorithm searches for an optimal position to recursively split an RNA sequence into two disjointed domains. In each iteration, the position is chosen to maximize the difference (measured by the earth mover′s distance (Yu et al., 2005)) between the distribution of interactions in the current domain and that in the two new sub-domains. This yields a hierarchical domain partition, and the top *k* domains in the hierarchy that minimize the maximal coefficient of variation of intra-domain interactions are selected as the output. This algorithm split the structure of the rRNA 18S into 4 domains and 28S into 6 domains, with the domain boundaries accurately demarcated for the two rRNAs (Supplementary Figure S4A and S4B).

We applied the method to define 23 structural domains for MR766 and 24 domains for PRVABC59 (Figure 3A and Supplementary Figure S2A). As expected, most PARIS interactions are clustered into intra-domain pairing structures. The enrichment ratio of intra-domain PARIS signal density over the background is about 32.3 for MR766 and 28.4 for PRVABC59 (Supplementary Figure S4C). The overall architectures of the two ZIKV genomic RNAs agree well, measured by the conservation of domain boundaries (13 boundaries are in common, p-value<0.001, permutation test. Supplementary Figure S4C).

Demarcating ZIKV genomes into small domains helps to accurately build their secondary structural models, as predictions have been successful for small RNAs especially when constrained with experimental probing data (Deigan et al., 2009, Wu et al., 2015). Here we used the software suite *RNAstructure* (Deigan et al., 2009) to predict the secondary structure for each domain separately, with icSHAPE scores as a constraint. We used the parameter that predicts the structure of 3′UTR most accurately for other domains (AUC=0.83 for PRVABC59 and AUC=0.80 for MR766). We verified the validity of our method by assessing the predictive performance on the 5′UTR (AUC=0.95 for PRVABC59 and AUC=0.92 for MR766). We also used a similar pipeline from a previous study of the HCV structures combined with a statistical tool R-scape to call covariant base pairs with 4,256 Flaviviridae virus genomes (Spitale et al., 2015, Rivas et al., 2017).

The method precisely reproduced the known secondary structural elements of the 5′-UTRs and 3′-UTRs within the ZIKV genomes (compare Figure 4, Figure S5 and Figure 2F). Most stems in these structural models demonstrate some covariation, suggesting that they are functionally conserved. In total, one RNA element in the 5′UTR domain (SLA) and four elements in the 3′UTR domain (SL1, SL2, DB2 and 3′SL) contain 71 co-variations in about 420 nucleotides (Figure 4). We did not find any covariation for one structural element, DB1 in the 3′UTR domain. Indeed, some flaviviruses, like YFV, lack DB1 (Villordo et al., 2016). The coding region, as expected, contains fewer evolutionary covariations (170 in about 10k nucleotides). However, the structural models do reveal some conserved structural elements in the coding region with covariations. For example, the capsid hairpin (cHP) in the coding region of the protein capsid contains 5 covariations. The cHP is involved in the translation and replication of flaviviruses, including DENV and WNV. Although the cHP hairpin structure is conserved, its sequence is quite flexible (Clyde et al., 2008). The functions of other structural elements remain to be elucidated (Figure 4). Overall, our structural analysis with icSHAPE and PARIS provided reliable structural models of two ZIKV genomes, and identified known and novel structural elements (Figure 4, Figure S5).

**Figure 4.**
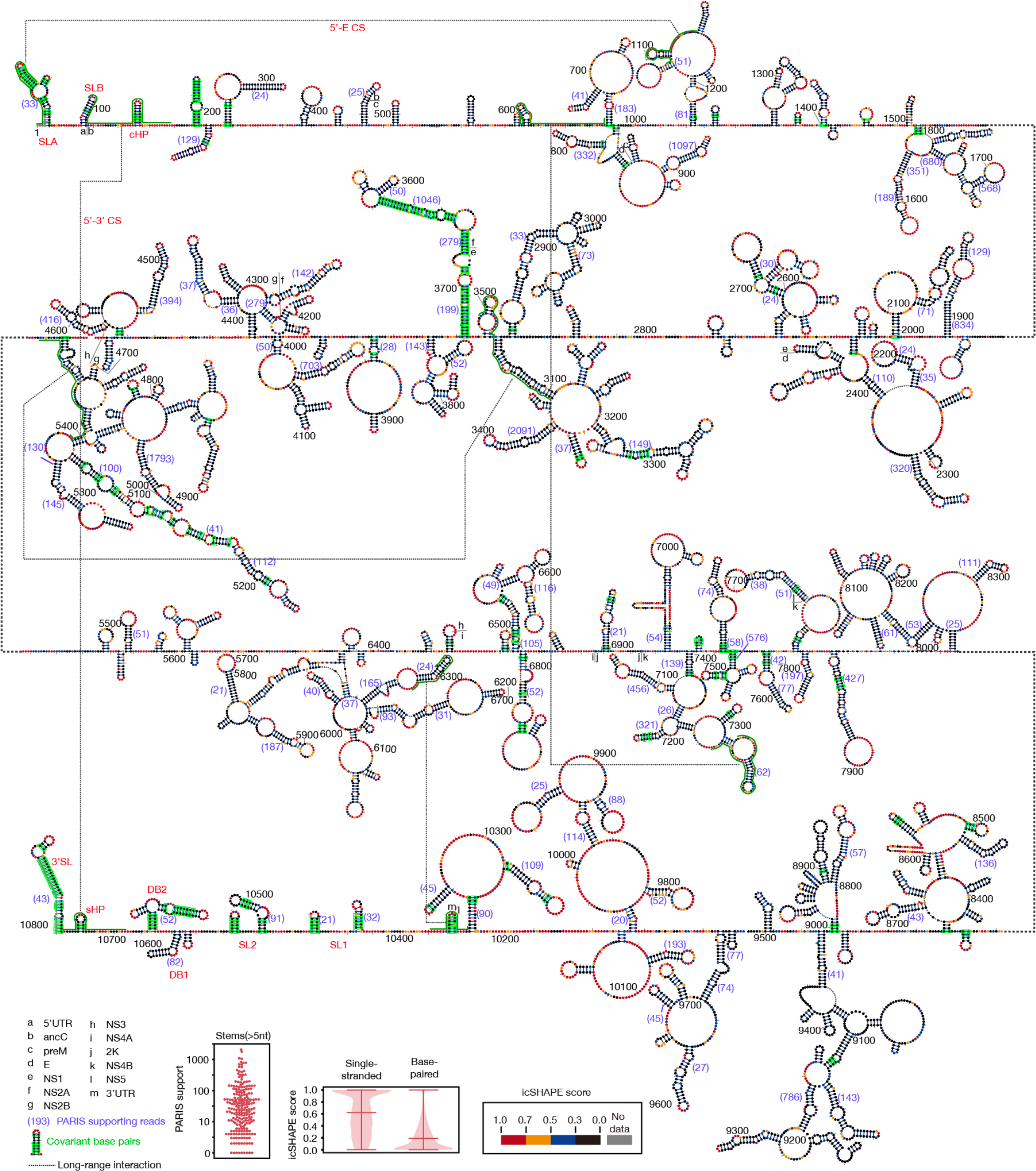
Full-Length structural model of the PRVABC59 RNA genome See also STAR Methods and Figure S5. Nucleotides are colored with icSHAPE reactivity scores. Base pairs with conserved covariation are boxed with green rectangle. Long range interaction in Figure 1D are indicated by dotted lines. The two boxplot insets at the bottom are the distributions of icSHAPE reactivity scores and PARIS supporting reads number for all interactions.

### A lineage-specific long-range interaction in epidemic ZIKV

The structural landscapes of the ZIKV genomic RNAs allowed us to investigate whether and how RNA structural elements may affect the infectivity of ZIKV. We hypothesized that lineage-specific structural elements might underlie outbreaks of epidemic strains. To search for lineage-specific structural elements, we first compared the structural flexibilities of the MR766 and the PRVABC59 strains from the icSHAPE data. We found 132 short regions that contain at least three nucleotides with substantially different icSHAPE scores (larger than 0.6, in a sliding window of 5 nucleotides). For each structurally distinct region, we defined it to be lineage-specific if the encoding sequence is conserved within each lineage but not in both. The analysis revealed a set of 68 elements (Supplementary Table S3); many of them could mediate lineage-specific RBP binding, RNA modification, etc. For example, the list contains a previously identified Musashi1(MSI1) protein binding site at the 3′-SL in the 3′UTR (Chavali et al., 2017). It has been reported that one base substitution in this site disrupts its binding to MSI1 in African strains (Chavali et al., 2017). Indeed, this substitution causes a structural change in MR766 and may render the binding with MSI1 energetically unfavorable.

Similarly, we defined lineage-specific intramolecular interactions from the PARIS results (271 short-range interactions for PRVABC59, 165 for MR766; and 127 long-range interactions for PRVABC59, 73 for MR766, Supplementary Table S4). Among these, we noticed a striking long-range interaction between the 5′ UTR (2-43nt) and the E protein coding region (1089-1134nt) in the PRVABC59 strain but not in the MR766 strain (Figure 5A). Although the 5′ UTR is well-conserved between the two strains, ten nucleotides in the E protein coding region are mutated to form this 5′ UTR-E interaction in PRVABC59, consisting of 37 base-pairs (Figure 5B and 5C). Phylogenetic analysis revealed strong conservation of these base pairs in all epidemic strains. In sharp contrast, this conservation is not observed in in pre-epidemic Asian strains nor any African strains. Interestingly, all the unconserved sites within the E protein coding region of pre-epidemic Asian strains and African strains are synonymous to maintain the same amino acid sequence at this region in the E protein of all strains (Figure 5B).

**Figure 5.**
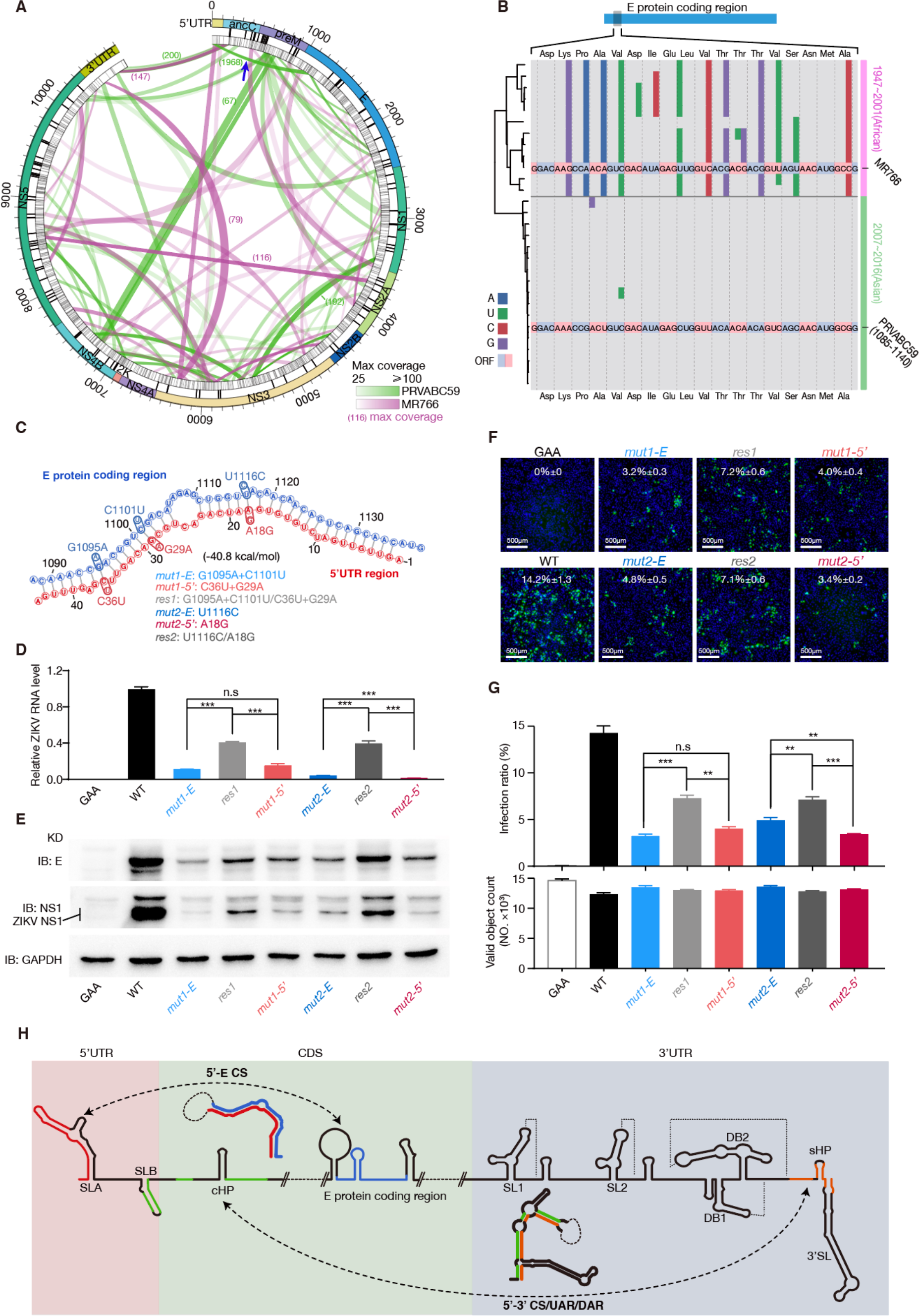
An Asian lineage-specific long-range intramolecular interaction may contribute to ZIKV infection. See also Table S4. A. Circos plot of the long-range interaction in the PRVABC59 genome and the MR766 genome. The outermost ring shows the genomic coordinates with protein annotation. The middle and the inner rings indicate amino acids and nucleotide sequence diversity, respectively. The color ribbons represent the long-range interactions of PRVABC59 (green) of MR766 (pink). The blue arrow points to the 5′-E CS interaction that only exists in PRVABC59. B. Nucleotide sequence diversity between African strains and Asian strains in the E protein coding region of the 5′-E CS. The two viral genomes are highlighted with PRVABC59 as the reference sequence. Mutations to the PRVABC59 genome are colored per type. Strains are organized according to the phylogenetic tree on the left. C. Predicted secondary structure model of the 5′-E CS interaction in PRVABC59, annotated with genomic coordinates. Blue circles represent designs of mutations and red circles represent rescues in the infection study. D. RT-qPCR quantitation of relative GZ01 viral RNA level from pellets of U87MG cells infected for 48h. The GAA mutant is a control with a defective NS5 RdRp domain (n=3). E. Western blotting of the ZIKV E and NS1 proteins from pellets of U87MG cells infected with GZ01 strains in 48h. (IB: immunoblotting). F. Immunofluorescence staining of ZIKV E protein in Vero cells infected for two days with GZ01 strains with equal viral titer as normalized by qRT-PCR. G. Summarized infection ratio as determined from (F) (up) and total number of cells (down). (n=3). H. Model of alternative ZIKV RNA genomic conformations including 5′UTR, E protein and 3′UTR local structures, long-range 5′-E CS and 5′-3′ CS interactions (dashed lines). Regions are marked to help identify pairing patterns. The p-value is calculated with student t-test. (*P<0.05, **P⁊<⁊0.01, ***P⁊<⁊0.001, unpaired two-tailed t-test). All error bars mean SEM for biological triplicates.

To investigate the function of this 5′UTR-E interaction in epidemic ZIKV, we used a full-length Asian strain GZ01, for which we have an infectious clone available for mutagenic analysis (Liu et al., 2017). The sequence of GZ01 is identical to that of PRVABC59 in the 5′UTR-E interaction region. We constructed mutants with synonymous mutations that disrupt base pairing without changing the E protein sequence in GZ01 (Figure 5C, G1095A+C1101A *mut1-E* or U1116C *mut2-E*). We also designed corresponding mutations in the 5′ UTR (Figure 5C, C36U+G29A *mut1-5′* or A18G *mut2-5′*). Finally, we engineered GZ01 mutants with the compensatory mutations in both the 5′ UTR and the E protein coding region to restore the base pairing (Figure 5C, G1095A+C1101A/C36U+G29A *res1* for *mut1-E* and *mut1-5′*, U1116C /A18G *res2* for *mut2-E* and *mut2-5′*). As a control, we included a non-replicating mutant ZIKV that lacks NS5 RNA dependent RNA polymerase function (“GAA”) (Liu et al., 2017). We transfected BHK21 cells with *in vitro* transcribed wild-type or mutant ZIKV RNA and harvested ZIKV particles in supernatant. To assess the full life cycle of ZIKV, the virions were generated from BHK cells transfected with viral genomic RNA, which was produced from *in vitro* transcription. The virions were quantitated with qPCR and equal titer of virions of each strain was used to infect U87MG cell line and infectivity was assessed by qPCR, Western blots and immunofluorescence. Similar to the GAA mutation, all of the mutations predicted to disrupt the long-range interaction (*mut1-E*, *mut1-5′*, *mut2-E* and *mut2-5′*) substantially reduced GZ01 infectivity. Importantly, the compensatory mutations *res1* and *res2* that would restore the 5′UTR-E interaction partially restored the infectivity in U87MG and Vero cells (Figure 5D-G). The fact that the combination of two deleterious mutations can rescue either mutant further demonstrate the reductions in infectivity by *mut1-E* and *mut2-E* are unlikely due to codon effects on protein translation. Instead, it strongly argues for the functional importance of the 5’UTR-E interaction.

The 5′ UTR of ZIKV RNA participates in the regulation of virus translation and replication through different pathways. It can initiate virus translation via a cap-dependent mechanism. For virus replication, the local structural element SLA functions as a promoter for the viral polymerase NS5. Another element SLB also facilitates replication by cyclizing the RNA genome via formation of a 5′-3′ complementary structure (5′-3′ CS) with the 3′ UTR (Ng et al., 2017) (Figure 5H). The switch between the translation and replication conformations is important for ZIKV infection (Liu et al., 2016). Our PARIS experiment identified a third conformation of the ZIKV 5′UTR, paring with the E protein coding region to form a 5′-E protein complementary structure (5′-E CS) (Figure 5H), which may play a role in virus replication or translation regulation.

## DISCUSSION

As an exogenous RNA in host cells, ZIKV genomes participate in many infection processes, presenting multiple levels of gene regulation. Studies have now started to reveal the complexity and significance of viral RNA function and regulation in RNA virus infection. For example, it has been shown that m^6^A modifications on the genomes of ZIKV and other flaviviruses modulate viral RNA metabolism in cells (Lichinchi et al., 2016, Gokhale et al., 2016). RNA structure can also reveal molecular properties and functions of viral RNAs (Mauger et al., 2015, Smola et al., 2015). Previous studies have uncovered RNA virus genome structures for HIV (Watts et al., 2009) and HCV (Mauger et al., 2015, Pirakitikulr et al., 2016). These studies identified many structural elements with known or uncharacterized regulatory functions, as well as global organizational principles that are important for viral processes. However, the studies are based on *in vitro* or *in viron* chemical probing, which do not completely reflect the complex regulation of RNA structures in infected cells. For ZIKV, due to the lack of global structural information, most studies have been focused on a limited set of RNA structures in the UTR regions. Here we report the first analysis that comprehensively revealed *in vivo* RNA structural regulation of ZIKV.

We combined two *in vivo* methods and obtained two complementary sets of ZIKV genome structural information with a high degree of agreement. Similar to the organization patterns of protein and DNA genome structures, we discovered structural domains in the ZIKV RNA genomes, with a large number of intra-domain interactions and a few inter-domain interactions. The domain organization was conserved between the two ZIKV genomes. Our data and previous studies showed that RNA secondary structures display more variability than RNA sequences (Mauger et al., 2015). Whether the domain organization is more conserved than simple RNA secondary structures, and its relevance to virus regulation, are interesting open questions.

We built structural models for each domain utilizing the *in vivo* probing data as constraints. The good agreement with existing, well-accepted models demonstrates the accuracy and reliability of our modeling algorithm. We did notice a few differences between our models and existing ones. For example, the SL1 structure at the 3′ UTR is slightly different in the two models (compare Figure 2F and Figure 4). However, this is possibly due to the *in vivo* and *in vitro* differences between RNA structures, as our experiments were performed *in vivo* and previous models combined information from *in silico* predictions and experiments including *in vitro* data. Our models of ZIKV viral genomes *in vivo* are more relevant to the virus life cycle in cells.

The structural models revealed both known and novel secondary structural elements within the ZIKV genomes. In particular, we noticed that some previously unknown long duplexes contain a large number of covariations, such as duplexes at the positions ~3600, ~7600 and ~8800 in the MR766 strain and at the position ~3600 in the PRVABC59 strain. In addition, the duplex at the position ~3600 is conserved in both strains, is predicted to be very stable, and has a large number of supporting reads, e.g., 125 PARIS reads in MR766 and 1046 reads in PRVABC59. The conservation and/or stability of these interactions suggests their possible functional significance.

We set out to use these RNA structural models to understand the differences in virus infectivity and pathogenicity between the African strains and the epidemic Asian strains. As an RNA virus with a high mutation rate, ZIKV RNA genomes have accumulated many nucleotide variations, some of which account for these functional differences. Conventional approaches evaluate protein differences (Yuan et al., 2017, Liu et al., 2017). Here, we focused on synonymous and non-coding mutations that change RNA structures without affecting protein sequences. We developed an analysis to identify lineage-specific RNA structural elements. We hypothesized that some of these lineage-specific structural elements could contribute to virus infectivity distinctions in different lineages and strains. We discovered an Asian lineage-specific long-range interaction between the 5′ UTR and E protein coding region, and confirmed its significance in ZIKV infection through mutagenesis and rescue experiments. The strong 5′UTR-E interaction might help ribosome binding and scanning on this 5′capped RNA genome or influence the SLA element in the 5′ UTR region and affect SLA-mediated recruitment of the Pol to the genome. The delicate balance between translation and replication regulation might alter the infectivity of the PRVABC59 strain. The detailed molecular mechanism of how this interaction contributes to ZIKV infection remains to be investigated. Nevertheless, the substantial changes in infectivity arising from a single nucleotide change exemplifies the important function of many structural elements in ZIKV genomes. Further characterization of other lineage-specific RNA structural elements may reveal additional RNA features that contribute to the viral infectivity and pathogenicity of epidemic strains.

Due to the lack of analytic tools, RNA structural studies have been mainly focusing on local structural elements. However, studies utilizing new techniques, including PARIS, SPLASH and LIGR-Seq, have started to reveal pervasive and dynamic long-range RNA interactions (Lu et al., 2016, Aw et al., 2016, Sharma et al., 2016). Some of these long-range interactions are conserved among different species (Lu et al., 2016). They can play essential roles in lncRNA modular organization and mRNA translation (Aw et al., 2016). Using PARIS, we for the first time systematically revealed many long-range interactions in ZIKV. We hypothesize that, in addition to the well-known 5′-3′ CS and our newly confirmed 5′-E CS, many of the other interactions we identified could be functional. Comparative analysis of the two ZIKV strains revealed that most long-range interactions in cells are not shared by the two species. In addition to the possibility of limitations in coverage, it is also possible that many of these are specific interactions that encode important functions for specific lineages and strains.

High-throughput technologies and system-wide analysis have started to unveil the structural landscape of RNA viruses and have provided a rich resource for the discovery of RNA structural elements that are important for virus infection (Watts et al., 2009, Mauger et al., 2015, Pirakitikulr et al., 2016, Ziv et al., 2018). RNA structures can function in a generic way, for example by affecting translation (Watts et al., 2009). They can also function in a specific way, exemplified by well-known structural elements at UTRs (Rouskin et al., 2014, Gebhard et al., 2011). A recent analysis that synonymously mutated blocks of the HIV-1 genome identified *cis*-acting elements that regulate splicing (Takata et al., 2018). In addition, it is well accepted that RNA structures influence protein and miRNA binding (Beaudoin et al., 2018, Taliaferro et al., 2016). It will be of interest to characterize the specific functions of the candidate structural elements discovered in this study, by large-scale mutational analysis. In sum, our approach and resource open a door to extensively investigate the function of RNA structures in virus infection.

## ACKNOWLEDGMENTS

We thank Xiaohua Shen (Tsinghua University) and Barry Honig (Columbia University) for helpful comments on the manuscript. This project is supported by the National the State Key Research Development Program of China (Grant No. 2016YFC1200300 to X.T.) and the National Natural Science Foundation of China (Grants No. 31671355, 91740204, and 31761163007 to Q.C.Z. and No. 31722030 to X.T.), the Beijing Advanced Innovation Center for Structural Biology to Q.C.Z, the Tsinghua-Peking Joint Center for Life Sciences and the National Thousand Young Talents Program of China to Q.C.Z. and X.T‥

## AUTHOR CONTRIBUTIONS

Q.C.Z. and X.T. conceived this project. P.L. analyzed all the results. Y.W. performed the icSHAPE experiments. L.T. performed the PARIS experiments. M.M. performed the virus mutagenesis and rescue studies with the help from L.S‥ J.Z implemented domain demarcating algorithm. Q.C.Z. and X.T. supervised the project. Q.C.Z., Y.W. and X.T. wrote the manuscript with inputs from all authors.

---

**Figure S1.**
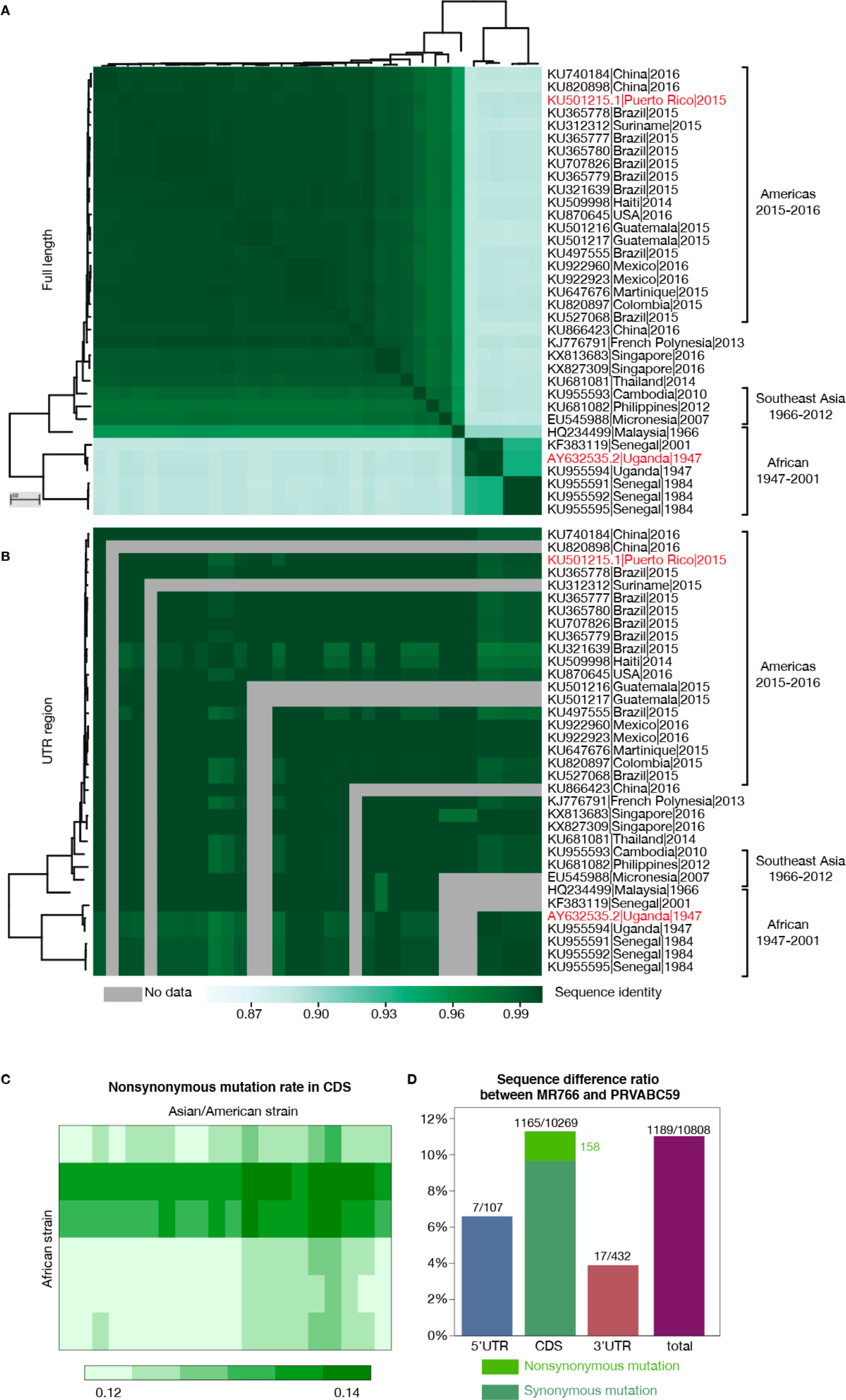

**Figure S2.**
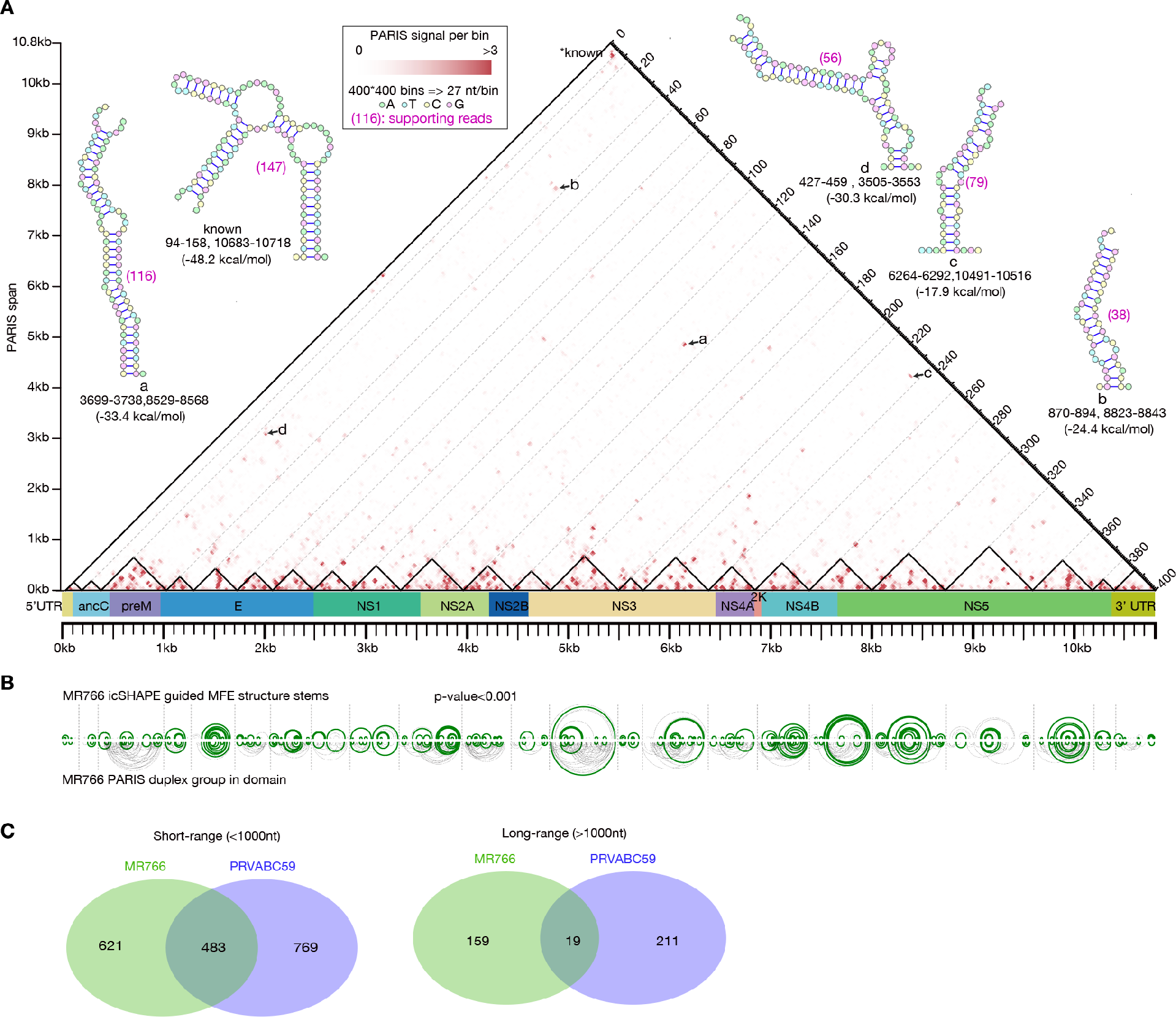

**Figure S3.**
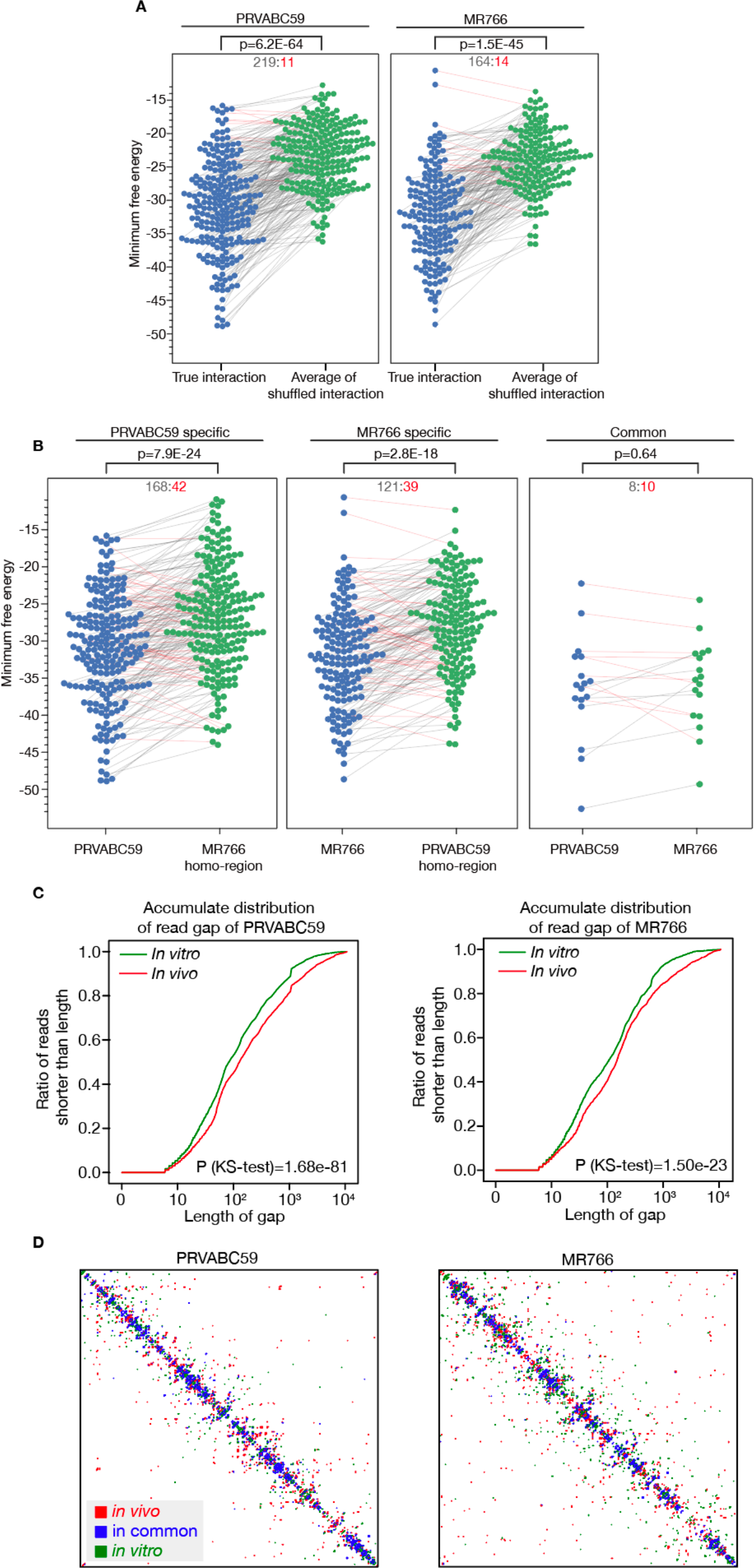

**Figure S4.**
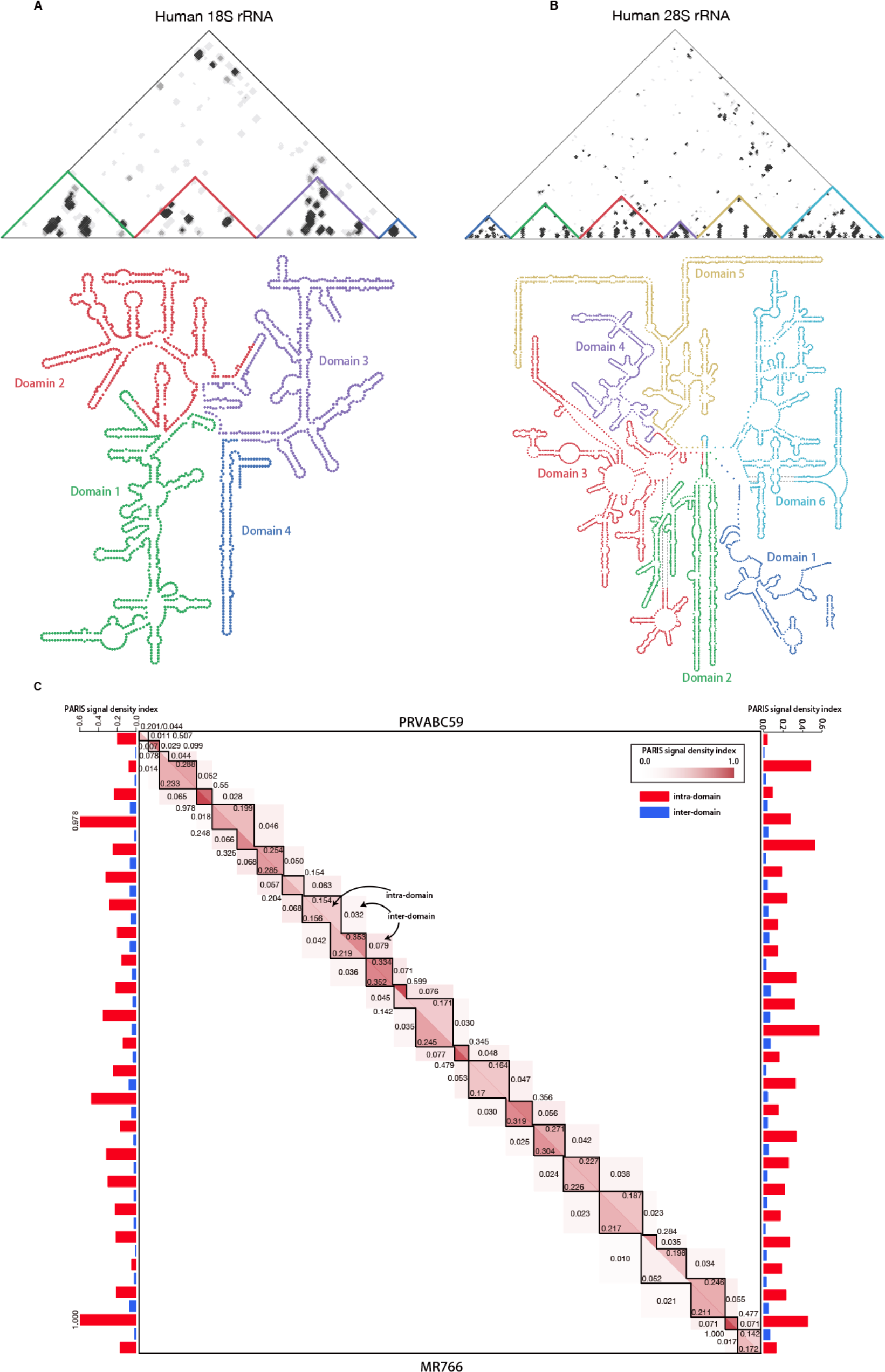

**Figure S5.**
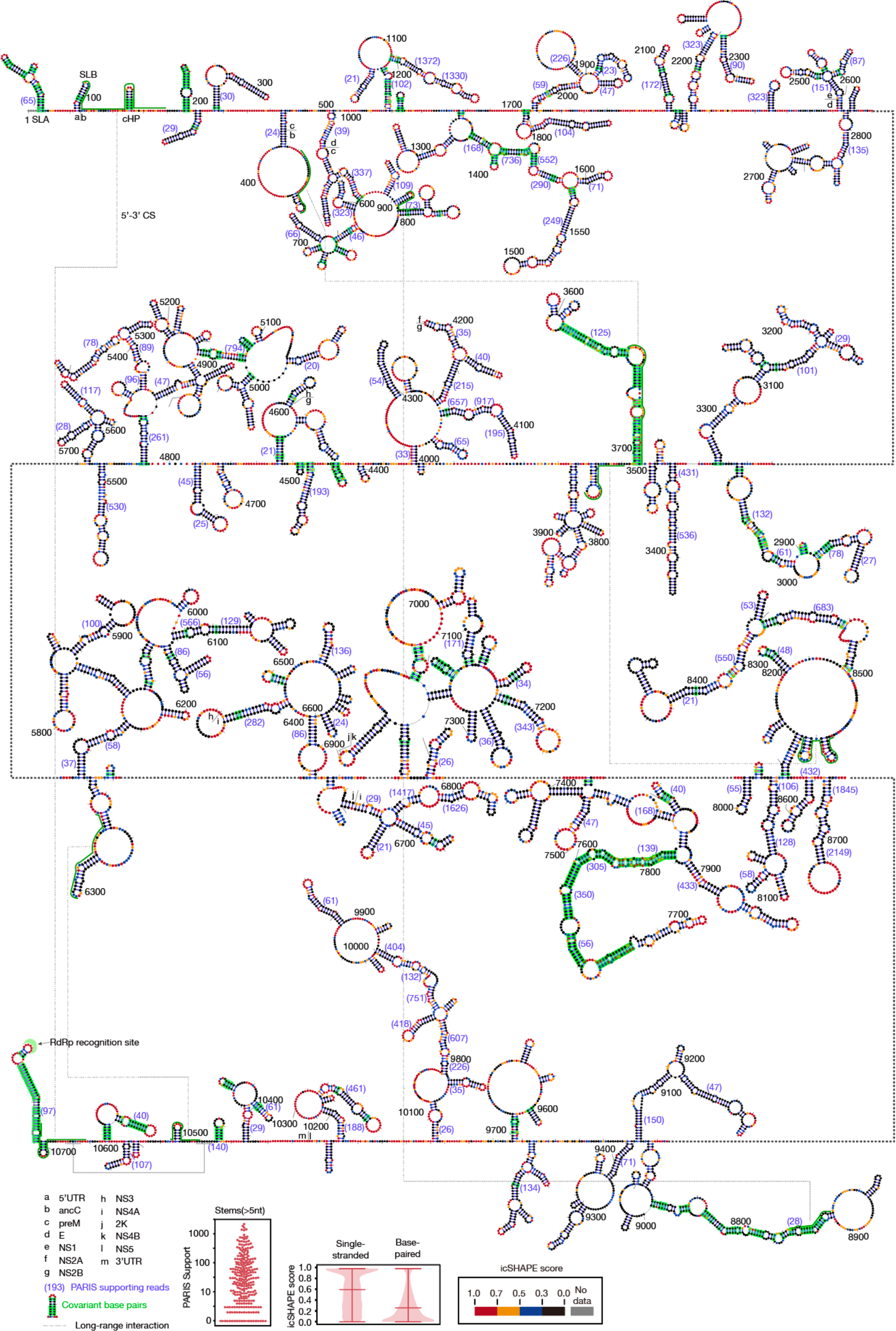

